# Expanding the DNA editing toolbox: novel lambda integrase variants targeting microalgal and human genome sequences

**DOI:** 10.1101/2023.09.22.559039

**Authors:** Siau Jia Wei, Asim Azhar Siddiqui, Lau Sze Yi, inivasaraghavan Kannan, Sabrina Peter, Zeng Yingying, Chandra Verma, Peter Droge, John F. Ghadessy

## Abstract

Recombinase enzymes are extremely efficient at integrating very large DNA fragments into target genomes. However, intrinsic sequence specificities curtail their use to DNA sequences with sufficient homology to endogenous target motifs. Extensive engineering is therefore required to broaden applicability and robustness. Here, we describe the directed evolution of novel lambda integrase variants capable of editing exogenous target sequences identified in the diatom *Phaeodactylum tricornutum* and the algae *Nannochloropsis oceanica*. These microorganisms hold great promise as conduits for green biomanufacturing and carbon sequestration. The evolved enzyme variants show >1000-fold switch in specificity towards the non-natural target sites when assayed *in vitro*. A single-copy target motif in the human genome with homology to the *Nannochloropsis oceanica* site can also be efficiently targeted using an engineered integrase, both *in vitro* and in human cells. The developed integrase variants represent useful additions to the DNA editing toolbox, with particular application for targeted genomic insertion of large DNA cargos.

## Introduction

Microalgae hold great promise as carbon-sequestering microbial cell factories for production of biofuels, nutraceuticals, pharmaceuticals, feed, and other biomolecules of commercial interest(1). Conventional strain engineering techniques to increase yields of endogenous biomolecules in microalgae have met with limited success, often due to deleterious trade-offs between productivity and cell-growth(2). Genome engineering is therefore a viable alternative to circumvent these issues and facilitate production of novel biomolecules by incorporating genes encoding exogenous enzymatic pathways(1). Targeted genome editing in microalgae via site-specific DNA cleavage has been shown utilizing clustered regularly interspaced short palindromic repeats (CRISPR/Cas9)(3), transcription activator-like effector nucleases (TALENS)(4), and zinc finger nucleases(5). Error-prone repair of lesions introduced results in corruption of native sequence context and perturbation of gene function. In the presence of exogenous donor DNA, homology-directed repair (HDR) can further result in targeted transgenesis(6, 7). To fully leverage algae’s potential as robust microbial cell factories, additional editing tools are required, particularly those facilitating targeted integration of even larger DNA cargos encoding complex multi-component pathways.

Bacteriophage and yeast derived site-specific recombinase (SSR) enzymes have enabled transgenesis of large (up to 133 kB) exogenous genes into the genomes of a diverse range of organisms(8, 9, 10, 11). SSR-mediated gene insertion does not rely on potentially error-prone endogenous DNA repair pathways, minimizing deleterious off-target effects commonly seen with other editing approaches(12, 13). However, intrinsic site-specificities of SSRs limits their applicability to targeting either cognate sequence motifs pre-introduced into the genome of interest or highly homologous “pseudo” sites. SSRs have therefore been engineered extensively to program novel sequence specificities and broaden their utility(10, 14, 15, 16, 17, 18, 19, 20).

λ integrase (Int) facilitates bi-directional transfer of the bacteriophage λ genome into the 21 bp *att*B target site in bacteria via specific recombination with the 241 bp *att*P attachment site resident in its genome(21, 22). Both integration and excision catalyzed by Int rely on bacterial DNA-bending factors (IHF, FIS and Xis) that bind to *att*P. Genomic editing in non-bacterial genomes is enabled by an engineered co-factor independent integrase variant (Int-h/218)(23, 24, 25, 26). Activity of Int-h/218 is also not compromised when used for *in vivo* recombination in *E.coli* (24).

Here, we describe engineering of λ integrase to target potential integration sites in the genomes of the diatom *Phaeodactylum tricornutum* and the algae *Nannochloropsis oceanica,* two microorganisms of significant interest for use as cell factories. The evolved integrase variants demonstrate a significant specificity switch towards these alternate sites, driven by mutation of residues both proximal and distal to bound DNA substrate. Using these engineered enzymes, we further demonstrate efficient targeted recombination into a single-copy sequence motif with homology to the *Nannochloropsis oceanica* integration site that is present in human cells.

## Materials and Methods

### Library generation and selection protocol

The integrase library was generated from C3-INT-HA pET22b(+) with GeneMorph II Random Mutagenesis kit using primers 8 and 9 (Table S1) and then re-amplified using primers tem-INTinf-ndeF and tem-INFinf-ecoR. The library was ligated into pINT selection vectors comprising *att*Phae2 and *att*Phae2Lsites after Nde1/EcoR1 restriction, followed by electroporation into TG1 cells and plated on selection plates containing increasing amounts of ampicillin (300 – 900 µg/ml). Plasmid DNA was isolated from colonies growing under the most stringent selection pressure (900 µg/ml ampicillin) and amplified with primers tem-INTinf-ndeF and tem-INFinf-ecoR. The library was ligated into the selection vector for the next round of selection under more stringent conditions (12mg/ml – 18 mg/ml). C4 was selected from the first round with 900 µg/ml ampicillin while C5 was selected from the second round with 18 mg/ml ampicillin.

### Recombination assay using *in vitro* translated proteins

Wild-type and variant integrase genes were amplified using INF-INT-nde1F and INF-INT-HA-ecoR1R (95°C 5mins, 95°C 5s, 55°C 20s, 72°C 1min, 25 cycles), cut with Nde1 andd EcoR1, and ligated into similarly cut pET22 vecor. Constructs were then amplified with primers 8 and 9 (95°C 5mins, 95°C 5s, 60°C 20s, 72°C 1min, 25 cycles). This created amplicons flanked with both T7 promoter and terminator sequences, which were used for *in vitro* transcription and translation (IVT). 20 ng of each integrase amplicon was used per reaction using PURExpress^®^ *In Vitro* Protein Synthesis Kit (NEB) in a total volume of 9 µl. Reactions were incubated at 30°C for 1 h. Intramolecular recombination was then assayed by adding 10 ng relevant plasmid substrate (pINT*att*B X *att*BL / pINT*att*Nanno X *att*NannoL */*pINT*att*HNanno X *att*HNannoL / pINT*att*Phae2 X *att*Phae2L) to a total volume of 10 µl. The mixture was allowed to incubate for 1.5 h at 30°C and 1µl of the reaction was taken for PCR with primers 1 and 8 (95°C 5mins, 95°C 5s, 55°C 20s, 72°C 1min, 30 cycles) or primers 3 and 4 (95°C 5mins, 95°C 5s, 60°C 20s, 72°C 30s, 30 cycles). The relative positions of primers used are shown in Figure 1. Sequences are shown in Table S1.

**Figure 1.**
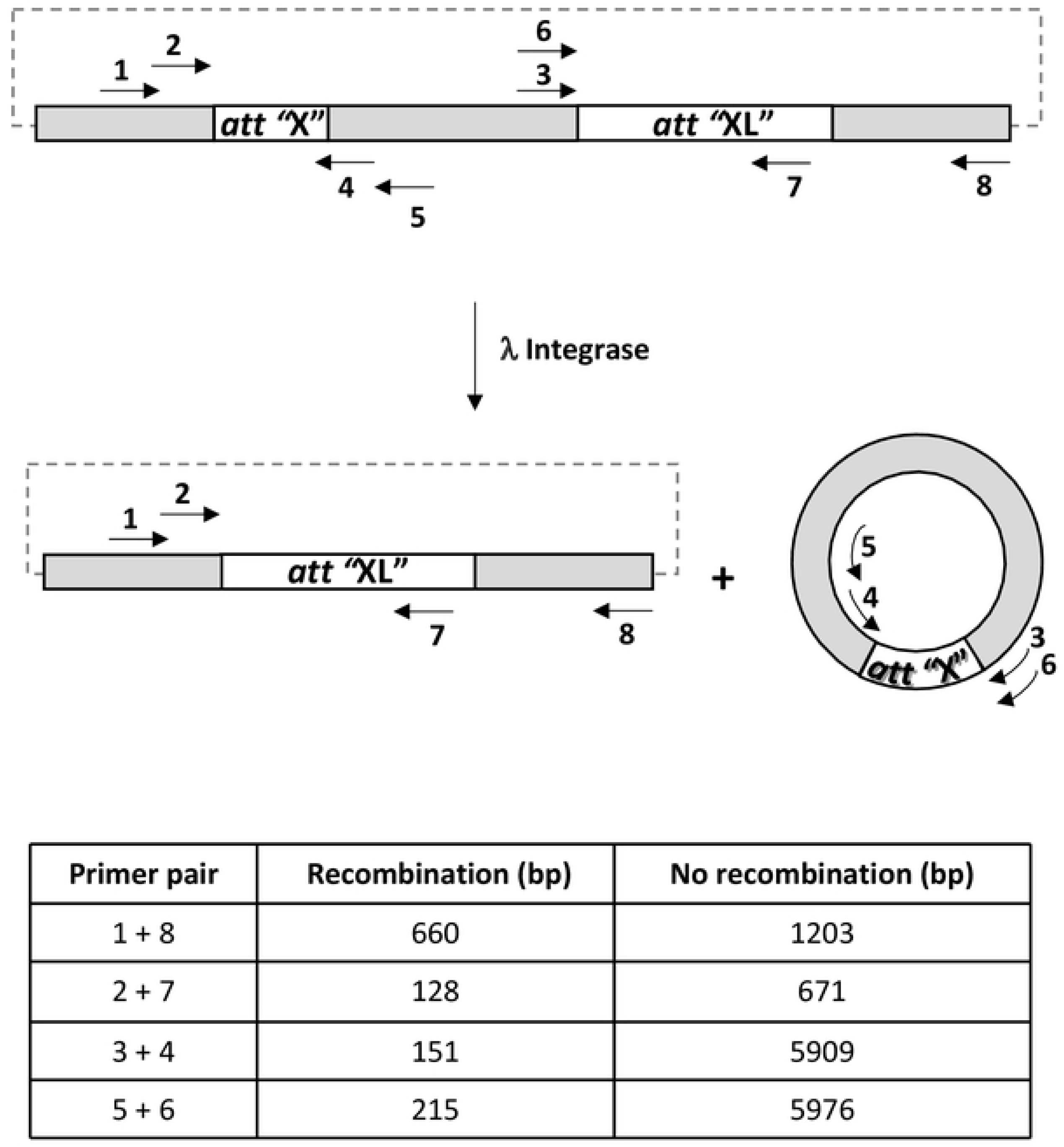
*In vitro* recombination assay. Int activity on plasmid DNA comprising appropriate *att* sequences yields the indicated products. Both the deletion event and resultant minicircle are detected by PCR using the indicated primers. Table shows expected size of amplification products for indicated primers pairs in presence or absence of recombination.

### Recombination assay using purified proteins

Recombinant integrase (17nM) was incubated with relevant plasmid substrate (1.3nM) (pINT*att*B X *att*BL, pINT*att*Nanno X *att*NannoL, pINT*att*HNanno X *att*HNannoL sites) in a total volume of 20µl in buffer comprising 150mM NaCl and 10mM Tris-HCl, pH7.8 at 37°C for 1 h. The mixture was diluted 1/10 using water and 1µl was used for real-time PCR quantification with 100nM of primers 5 and 6. End-point PCR was carried out by taking 1µl of undiluted mixture for amplification with primers 1 and 8.

### Cell culture

HT1080 cell line was maintained in Dulbecco’s Modified Eagle Medium (DMEM) growth medium supplemented with 10% FBS, 1% L-glutamine and 100 Units/ml of Penicillin and Streptomycin each (Gibco, Life technologies) at 37°C under 5% CO_2_ in humidified condition. For selection of puromycin-resistant recombinants, puromycin (Gibco, Life technologies) was added to the growth medium (1 μg/ml final concentration). Trypsin-EDTA (Gibco, Life technologies) was used for detaching the adherent cells for passaging.

### Transfection

For transfections in HT1080, 3 × 10^5^ cells were seeded per well of a six-well plate (TPP, Switzerland) in DMEM growth medium a day before transfection to obtain 70–90% confluence. Transfections were carried out with Lipofectamine 2000 (Invitrogen, Life technologies) with DNA to Lipofectamine2000 ratio of 1 μg:3 μl. For every transfection per well, DNA (1 µg targeting plasmid, 0.5 µg scIHF2 expression vector with or without 0.5 µg Int-C7 plasmid) and Lipofectamine 2000 were incubated separately in 50 µl of Opti-MEM medium (Life Technologies). The lipid complexes were prepared by mixing DNA and Lipofectamine 2000 reagent and incubating for 20 min at room temperature. The transfection mix was added dropwise onto the cells (under DMEM growth medium without antibiotics) and transfection was allowed to proceed overnight before cells were transferred to a 10 cm cell culture plate (TPP, Switzerland).

### Antibiotic selection and screening for targeted cell clones

Forty-eight hours post-transfection, selection with puromycin in growth medium at the concentrations indicated above was initiated. Selection medium was replaced once in 2 days until colonies expanded to ∼0.3–0.4 cm in diameter. At this stage, the colonies were picked by carefully scraping patches of cells with a pipette tip and transferred to 24-well plates for clonal expansion. The clones were sequentially expanded from 24 wells to six-well plates. Genomic DNA was extracted using DNeasy Blood & Tissue Kit (Qiagen, GmbH) as per manufacturer’s protocol for PCR screening.

### Identification of recombination events by PCR screening

PCR was performed using GoTaq Flexi DNA polymerase (Promega) to amplify both junctions using primers listed in Table 1 and routinely 500 ng of genomic DNA from parental cells or each recombinant clone as template in 25 μl reactions. The thermal cycling parameters used for primary PCRs from transfected and puromycin selected parental cells were as follows: initial denaturation at 95°C for 2 min, 35 cycles of denaturation at 95°C for 1 min, annealing at 57°C for 1 min and extension at 72°C for 1 min 10 sec (left junction) and 1 min 40 sec (right junction), and a final step of 72°C for 5 min. This was followed by a nested PCR with 1 ul of template from the product of primary PCR as follows: 95°C for 2 min, 35 cycles of denaturation at 95°C for 30 sec, annealing at 57°C for 30 sec and extension at 72°C for 20 sec (left junction) and 1 min (right junction), and a final step of 72°C for 5 min. The PCR samples were analyzed by electrophoresis in 1% agarose gels in 0.5× TBE (Tris-boric acid-EDTA buffer) containing 0.5 μg/ml ethidium bromide, and PCR-amplified products were compared with DNA standard markers and digitally documented under UV illumination (Quantum Vilber Lourmat, Germany). PCR-amplified products were analyzed by sequencing.

**Table 1.**
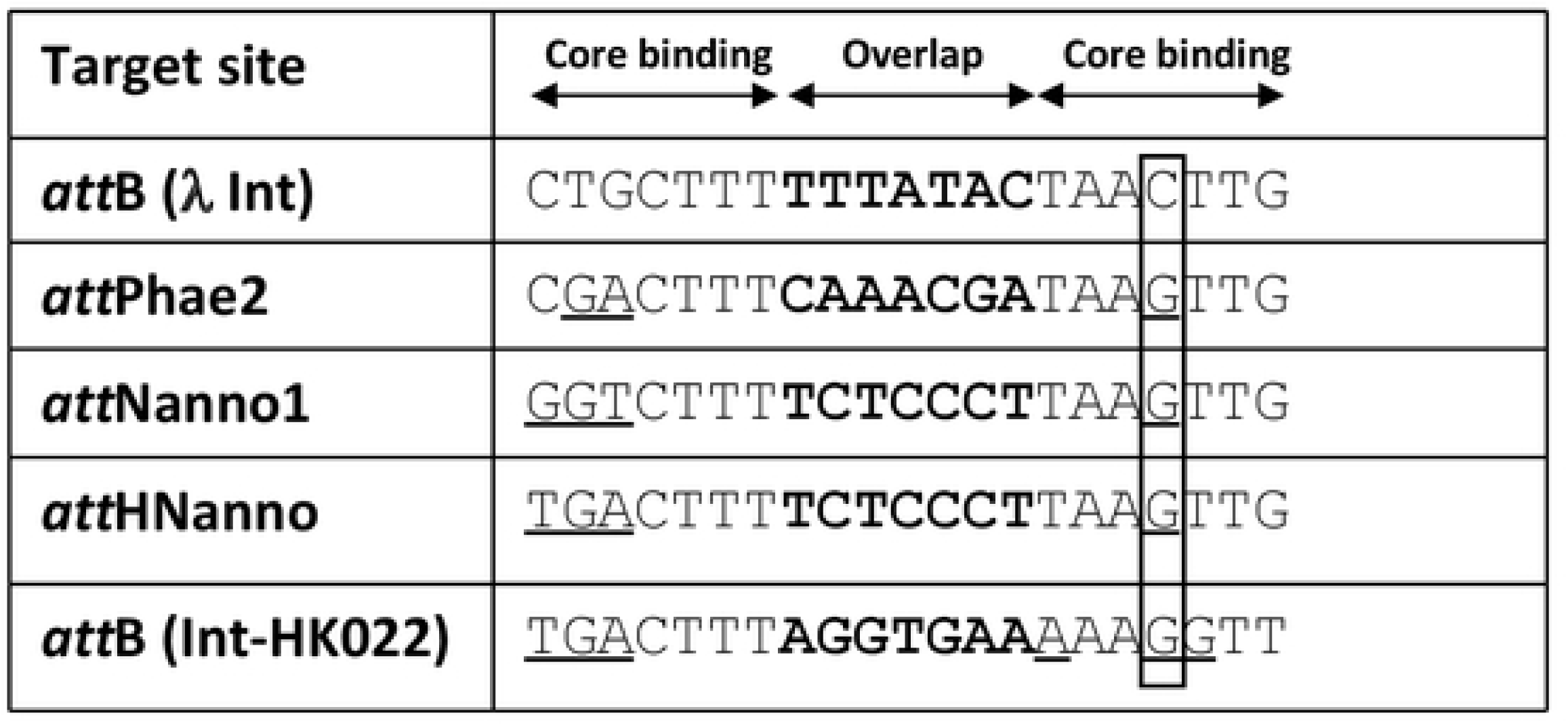
Target site DNA sequences used in the study. The overlap sequences are in bold. Sequence deviations from the endogenous λ integrase *att*B sequence outside the overlap region are underlined. The boxed region denotes key cytosine to guanine change important for specificity determination.

### Molecular Dynamics simulations

The available crystal structure (pdb: 1Z1G(27)) of lambda integrase tetramer bound to a DNA holiday junction (excluding the integrase residues 1 to 74 and mutating the Phe342 back to wildtype Tyr342) was used to generate several mutant models of lambda integrase – DNA complexes. Both the wildtype (WT) and mutant models were subject to Molecular Dynamics (MD) simulations with the pmemd.CUDA module of the program Amber18(28). All atom versions of the Amber force fields FF14SB(29) and FF99BSC0(30) were used to model the protein and DNA respectively. The *Xleap* module was used to prepare the system for the MD simulations. Each simulation system was neutralized with an appropriate number of counterions. Each neutralized system was solvated in an octahedral box with TIP3P(31) water molecules, with at least a 10 Å boundary between the solute atoms and the borders of the box. During the simulations, Lennard Jones and short-range electrostatic interactions were treated using a cut-off scheme and the long-range electrostatic interactions were treated with the particle mesh Ewald method [6] using a real space cut-off distance of 9 Å. The Settle(32) algorithm was used to constrain bond vibrations involving hydrogen atoms which allowed a time step of 2 fs to be used during the simulations. Solvent molecules and counterions were initially relaxed using energy minimization with restraints on the protein and DNA. This was followed by unrestrained energy minimization to remove any steric clashes. Subsequently the system was gradually heated from 0 to 300 K using MD simulations with positional restraints (force constant: 50 kcal mol-1 Å-2) on the protein and DNA over a period of 0.25 ns allowing water molecules and ions to move freely. During an additional 0.25 ns, the positional restraints were gradually reduced followed by a 2 ns unrestrained MD simulation to equilibrate all the atoms. For each system, a 250 ns production MD at 300 K was carried out in triplicate (assigning different initial velocities to propagate each MD simulation). To enhance the conformational sampling, the systems were subjected to accelerated MD (aMD)(33) simulations as implemented in AMBER 18(28). aMD simulations were performed on all three systems using the “dual-boost” version(34). Conventional MD simulations mentioned earlier were used to derive the aMD parameters (EthreshP, alphaP, EthreshD, alphaD). aMD simulations were carried out for 500 ns each. Simulation trajectories were visualized using VMD(35) and figures were generated using Pymol(36).

## Results

### Selection of **λ** integrase variants with altered specificities

Potential λ integrase targeting sites in the genome of the marine diatom *Phaeodactylum tricornutum* were identified *in silico* using the canonical 21 bp bacterial *att*B *sequence* (Table 1) to query sequence databases. A highly similar site, termed *att*Phae2, was identified with deviations from *att*B at 3 nucleotide positions in the core binding motifs flanking the highly variable overlap motif where reciprocal DNA strand exchange occurs (Table1). Recombination into *att*Phae2 was assessed using an *in vitro* plasmid-based assay, whereby intramolecular recombination between *att*Phae2 and *att*Phae2L (a modified *att*L sequence adjusted for the overlap and 5’region found in *att*Phae2) was measured by real-time PCR (Figure 1). The parental integrase (Int-h/218) showed minimal activity for attPhae2 x attPhae2L recombination. In contrast, the previously described hypermorphic Int C3 variant(10) showed ∼22-fold improved recombination of this substrate pair (Figure 2A). Int C3 was therefore used as starting point for further engineering using an *in vivo* directed evolution platform that couples correct recombination of plasmid-borne substrate pairs to survival of *E. coli* on selective media(10). Random mutagenesis of Int C3 followed by 2 rounds of selection yielded two hits, C4 and C5 (Table 2) with ∼ 2-fold improved activity over Int C3 for recombination using *att*Phae2 x *att*Phae2L DNA substrates (Figure 2A).

**Figure 2.**
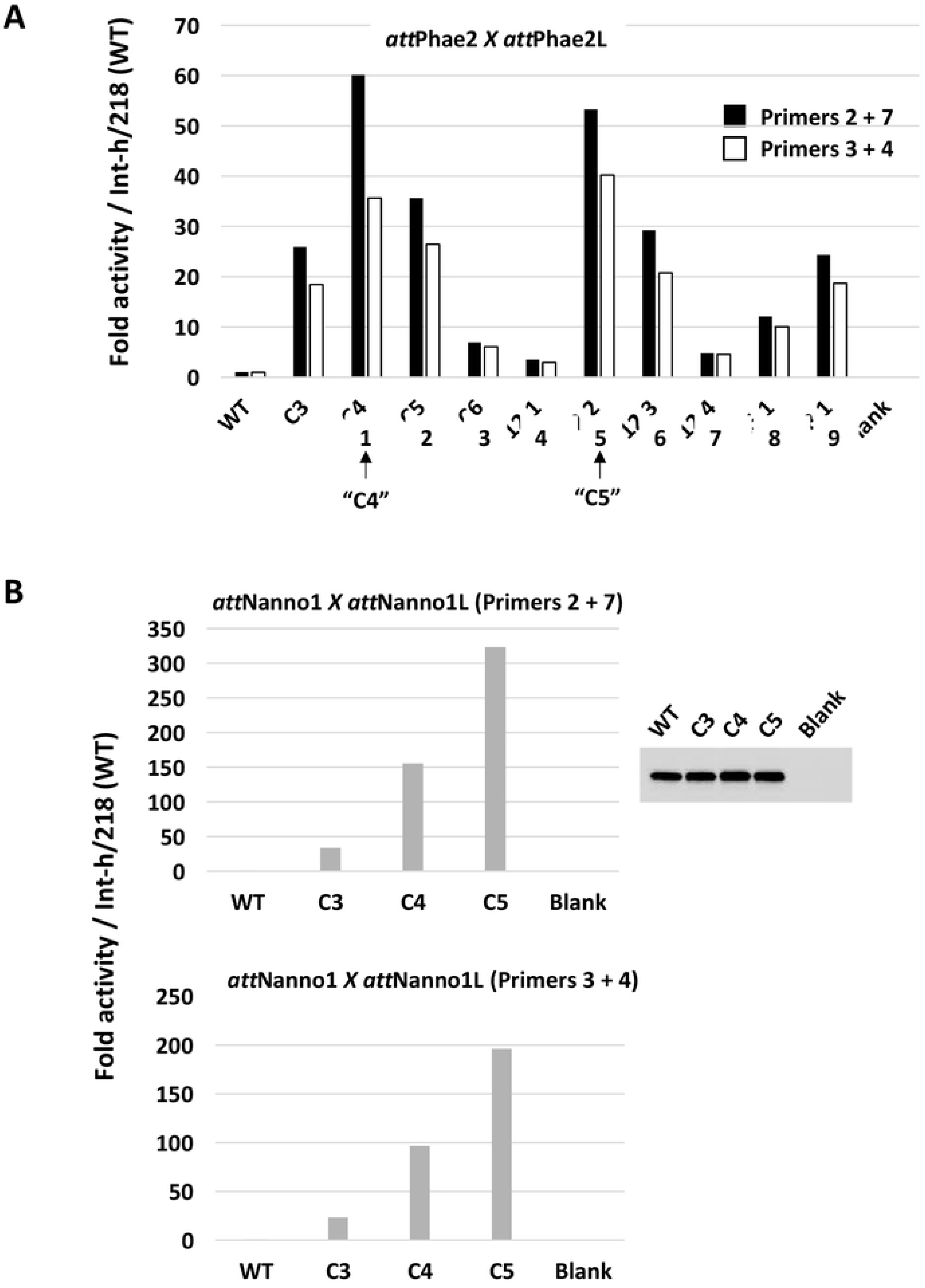
Recombination activity of selectants measured by qPCR. **A)** *In vitro* translated integrases were incubated with *att*Phae2 x *att*Phae2L plasmid substrate and recombination assessed by real-time PCR using primers 2 + 7 (black bars) or primers 3 + 4 (white bars). Activity is presented as fold increases over Int-h/218 (WT). Selectants 1 and 5 showing improved activity over C3 are respectively renamed Int C4 and C5. Blank corresponds to addition of *in vitro* translation extract only (no integrase expressed). **B)** As in **A** using indicated integrase variants and *att*Nanno1 X *att*Nanno1L plasmid substrate. Top graph uses primer 2 + 7 pair. Bottom graph uses primer 3 + 4 pair. Inset shows expression levels of integrases by Western blot. NTC: *in vitro* translation mix only (no integrase expressed).

**Table 2.** Mutations present in previously disclosed integrase variants (Int-h/218 and C3) and novel variants identified in this study (C4 – C7).

| Variant | Mutations |
| --- | --- |
| Int-h/218 | E174K, E218K |
| C3 | E174K, E218K, I43F, E319G, D336V |
| C4 | E174K, E218K, I43F, E319G, D336V, **P14L, D71V, N99D, E277V** |
| C5 | E174K, E218K, I43F, E319G, D336V, **H329R** |
| C6 | E174K, E218K, I43F, E319G, D336V, **P14L, D71V, N99D, E277V, H329R** |
| C7 | E174K, E218K, I43F, E319G, D336V, **N99D, H329R** |

Int C4 and C5 were subsequently assayed for recombination of a putative target site identified in the genome of the microalgae *Nannochloropsis oceanica* (*att*Nanno1), differing from *att*Phae2 by two and six bases in the left arm of the core binding and overlap sequences respectively (Table 1). At similar expression levels, both showed notably improved activity over Int C3 for recombination of *att*Nanno1 x *att*Nanno1L (∼5 and 10-fold respectively) (Figure 2B). These were tested again along with a further variant termed C6, comprising all the mutations present in C4 in addition to the H329R mutation identified in C5 (Table 2). As before, all variants showed improved activity over C3 for recombination of *att*Nanno1 substrates (Figure 3). C6 was more active than either C4 and C5, highlighting an additive effect of combining mutations from C4 and C5. Notably, the C4 and C6 variants recombined endogenous *att*B x *att*BL with reduced efficiency compared to C3 and C4, indicated by lower yield of the 660 bp PCR product scoring for correct recombination (Figure 3B). Analysis of mutations in C4 and C6 highlighted N99D as potentially responsible for the observed specificity switch phenotype based on the proximity of N99 to the DNA substrate in the crystal structure and prior reports highlighting this residue as a specificity determinant (37, 38, 39). We therefore generated a panel of constructs to interrogate the N99D mutation. These comprised Int-h/218 + N99D, Int C3 + N99D, Int C4 + D99N reversion, and Int C6 + D99N reversion. When tested using *att*Nanno x *att*NannoL substrates, recombination by Int-h/218 + N99D and Int C3 + N99D was improved over Int-h/218 and Int C3 (Figure 4). Conversely, the D99N reversion in Int C4 and Int C6 clearly led to reduced efficiency of recombination. These results highlight a key role of the N99D mutation for the observed specificity switch of the C4 and C6 Int variants.

**Figure 3.**
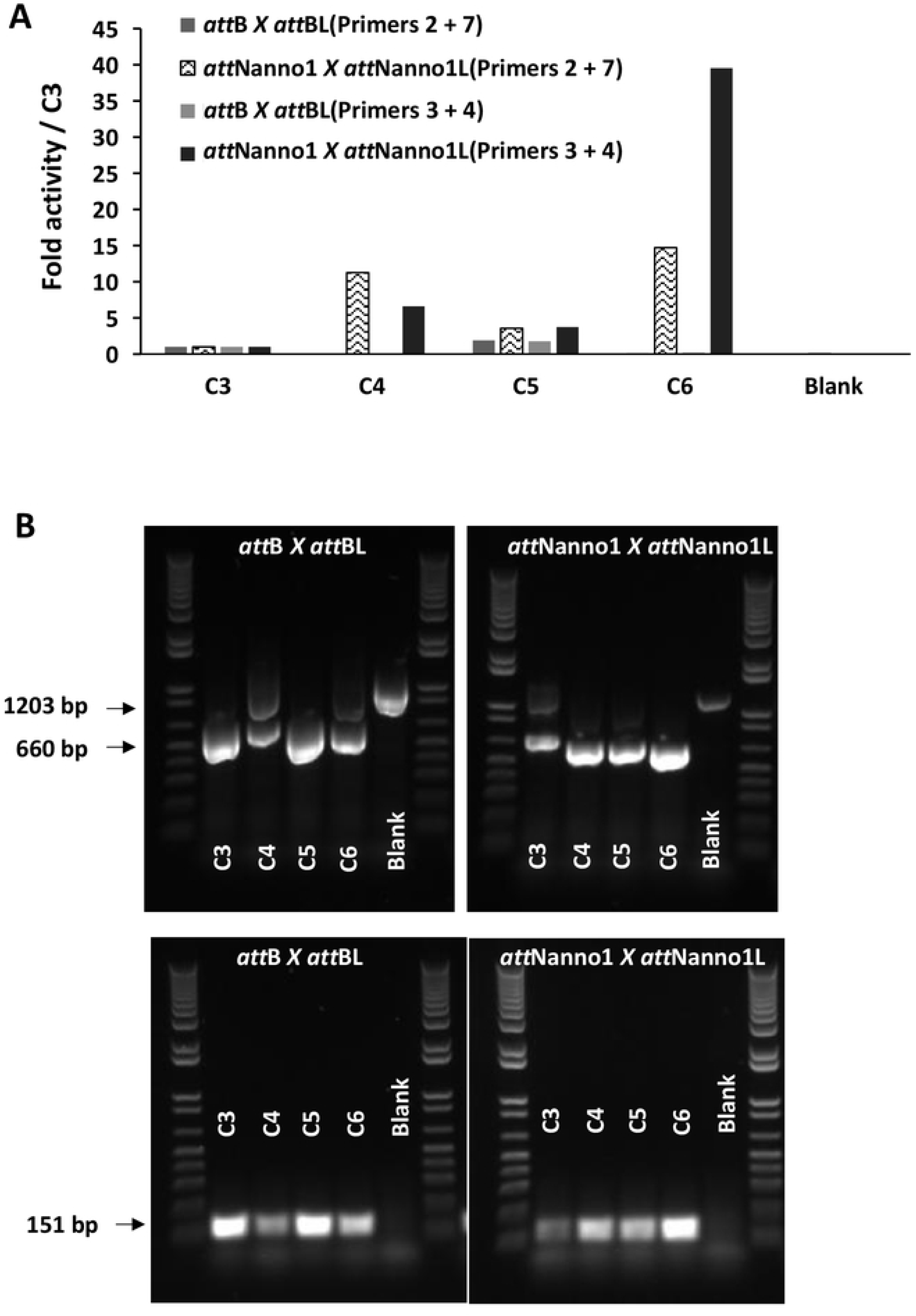
Recombination activity of selectants. **A)** *In vitro* translated integrases were incubated with indicated DNA substrates and activity determined by real-time PCR using either primers 2 + 7 or primers 3 + 4. Blank corresponds to addition of *in vitro* translation extract only (no integrase expressed). **B)** Same as A, except that post incubation an aliquot of each reaction was used in end-point PCR using primer 1 + 8 pair (top panels) or primer 3 + 4 pair (bottom panels). Using the first primer pair, successful recombination of the plasmid substrate yields a 660 bp fragment (lower arrow), whilst non-recombined plasmid yields a 1203 bp fragment (upper arrow). Recombination measured by the primer 3 + 4 pair yields a 151 bp band (arrowed).

**Figure 4.**
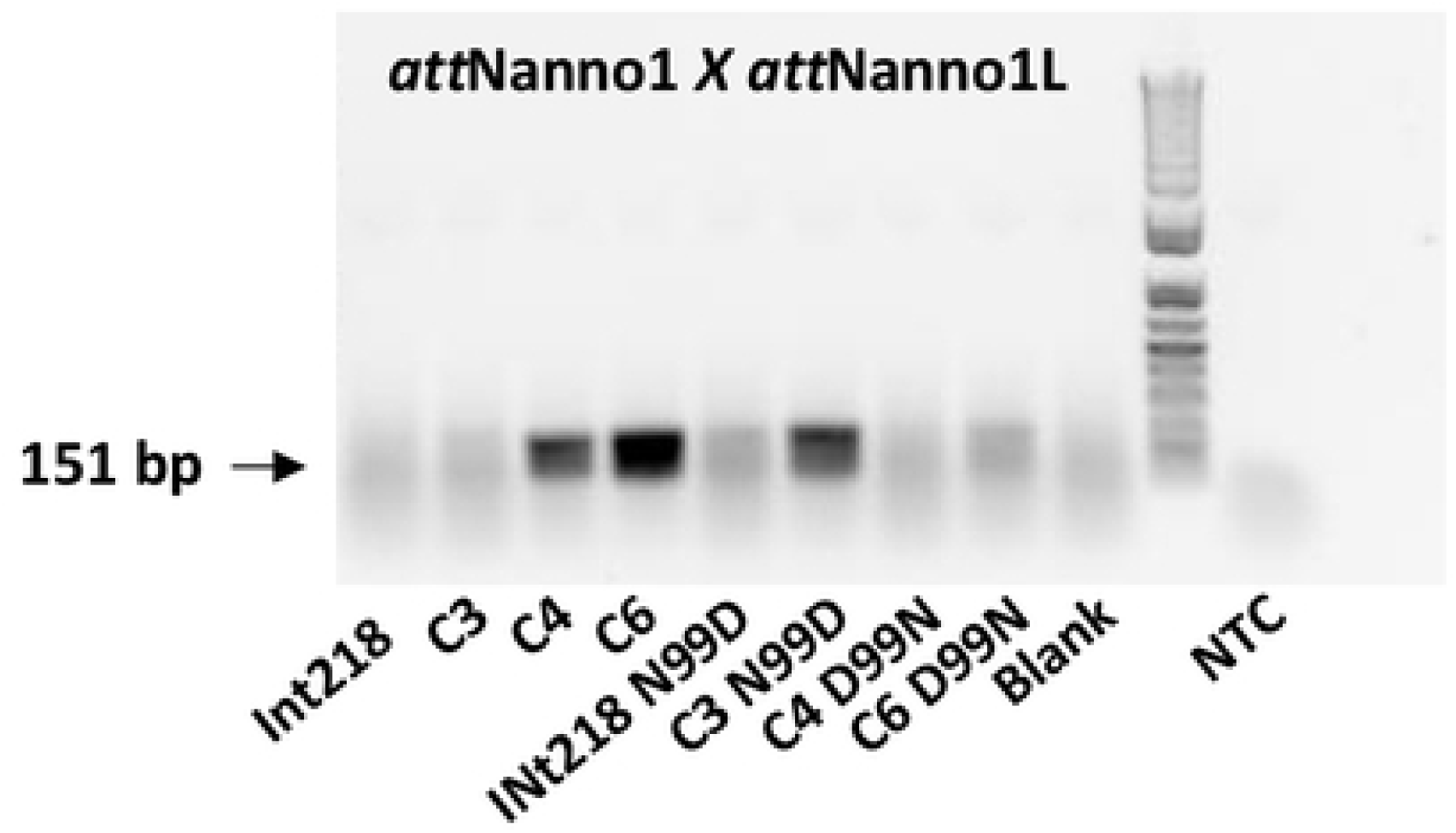
N99 acts as a specificity determinant. The indicated integrases were made using *in vitro* translation and incubated with *att*Nanno1 x *att*Nanno1L plasmid substrate. Scoring of recombination post incubation was carried out by end-point PCR using the primer 3 + 4 pair. Recombination measured by these primers yields a 151 bp band (arrowed). Repeat experiment shown in Figure S1.

We next tested recombinantly expressed and purified Int C3, C4 and C6 protein variants for more quantitative evaluation of function using a real-time PCR assay. The confirmatory results in Figure 5 show enhanced recombination of *att*Nanno1 substrates by C4 (∼11 fold improved over C3) and C6 (∼32 fold improved over C3). We also evaluated recombination of a sequence identified in the human genome with homology to *att*Nanno1 termed *att*HNanno (Table1). Int C4 showed ∼106-fold improved recombination of HTN over Int C3. C6 showed ∼369 fold improved recombination of *att*HNanno over C3. As before, recombination of *att*B x *att*BL substrate was less efficient for C4 and C6 compared to C3. We further confirmed these results using purified proteins by end-point PCR assay (Figure 6).

**Figure 5.**
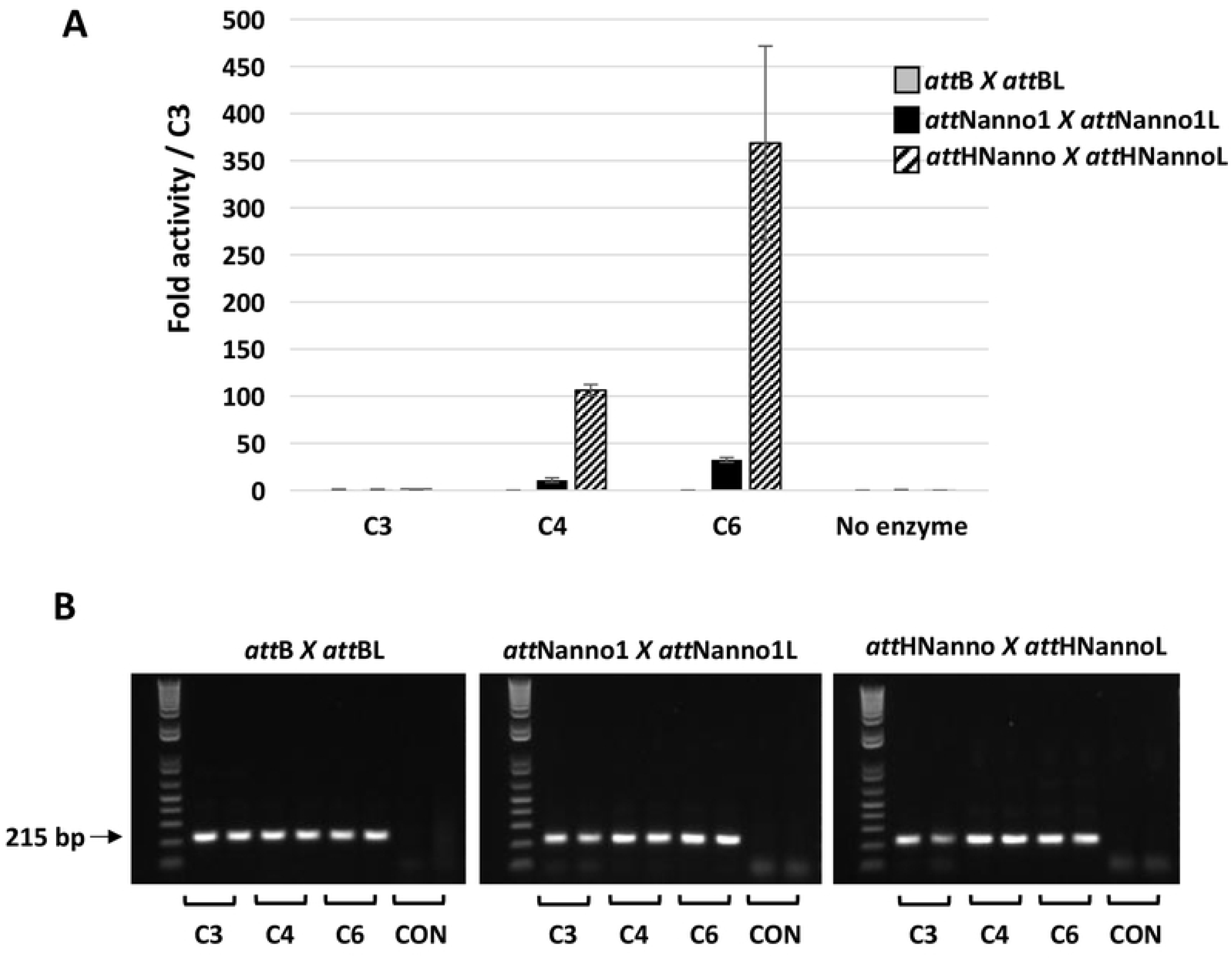
Recombination activity of selectants using purified proteins. **A)** Purified integrases (C3, C4, C5) were incubated with indicated plasmid DNA substrates and activity determined by real-time PCR using the primer 5 + 6 pair. n=2 ± SD. **B)** The real-time PCR reactions products analysed on agarose gel. Arrow indicates size of correct band (215 bp) indicating recombination event. CON: no enzyme in recombination reaction. Repeat end-point gel shown in Figure S2.

**Figure 6.**
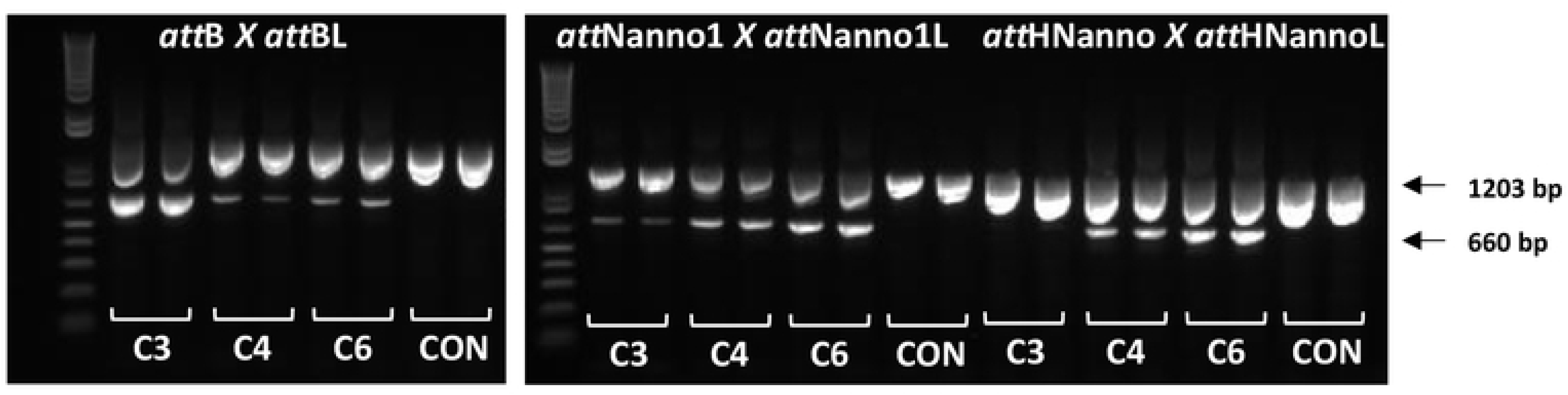
Recombination activity of selectants using purified proteins. Purified integrases (C3, C4, C6) were incubated with indicated plasmid DNA substrates and activity determined by end-point PCR using primers 1 and 8. The PCR products corresponding to unrecombined (1203 bp) and recombined (660 bp) template DNA are indicated by arrows. CON: no enzyme in recombination reaction. Duplicate results are shown.

Based on results highlighting a crucial role for N99D mutation in observed phenotype we generated a construct termed Int C7 (Table 2) comprising the expected minimal set of mutations required for improved recombination of the novel *att*Nanno1 and *att*HNanno substrates (N99D and H329R) in addition to the core C3 mutations. Expressed and purified C7 showed ∼412-fold improvement over C3 for recombination of *att*Nanno1 substrates and ∼1347-fold improvement for recombination of HTN substrates (Figure 7A). The respective values for C6 were ∼376 and ∼483. C7 was ∼1.1 and ∼2.8-fold more efficient than C6 for respective recombination of *att*Nanno1 and *att*HNanno recombination targets. Conversely, C6 and C7 respectively recombined *att*B x *att*BL ∼14 and 6.8-fold less efficiently than C3 (Figure 7B). This represents an ∼5264-fold change in specificity for C6 recombination of *att*Nanno1 versus *att*B substrates and an ∼6762-fold change in specificity for recombination of *att*HNanno substrates. The corresponding values for C7 are ∼2801 and ∼9159 respectively. Repetition of this experiment using a 10-fold lower final concentration of purified integrase and DNA substrate (Figure 8) led to similar results, with an ∼1241-fold change in specificity for C6 recombination of *att*Nanno1 versus attB substrates and an ∼1139-fold change in specificity for recombination of *att*HNanno substrates. The corresponding values for C7 were ∼597 and ∼1146 respectively.

**Figure 7.**
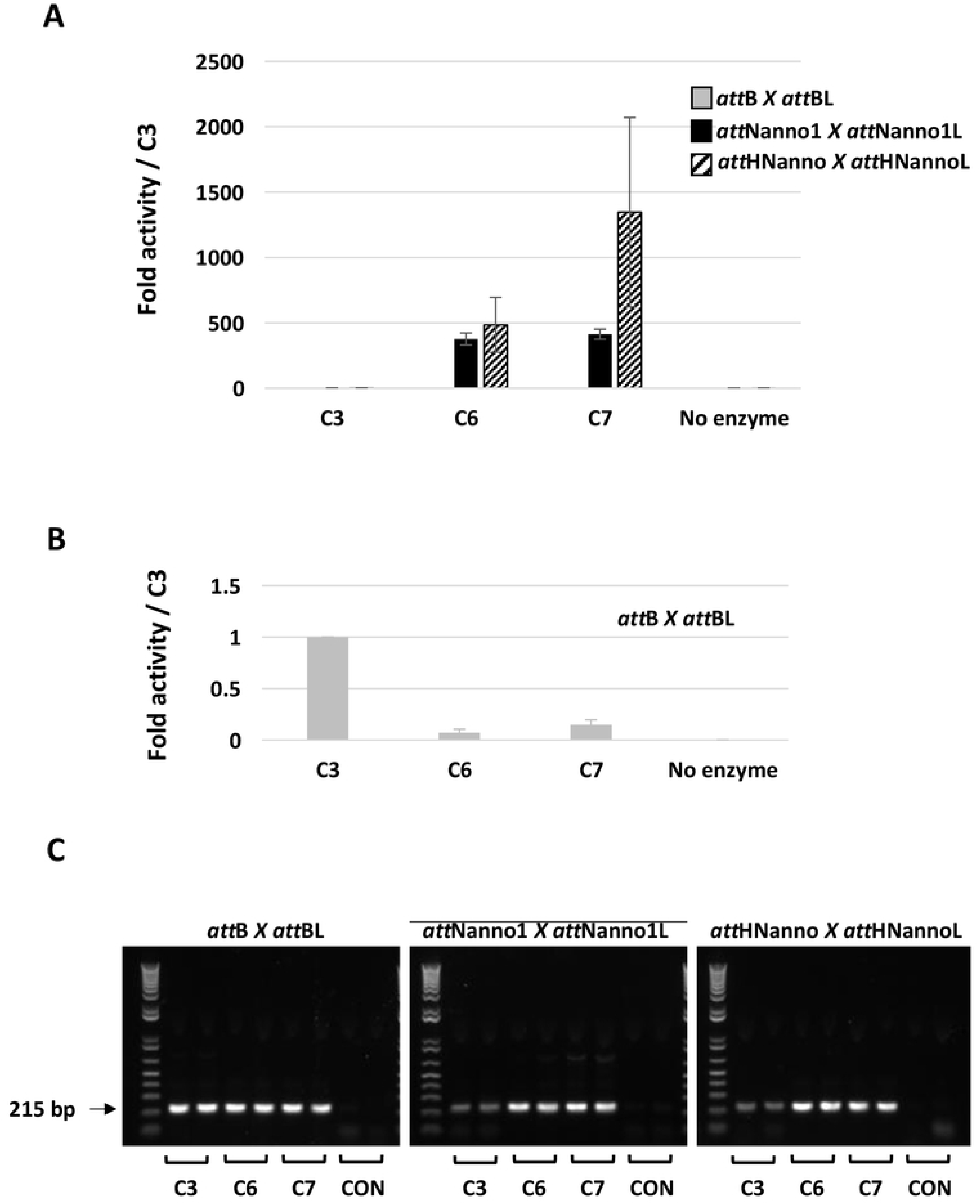
Recombination activity of selectants using purified proteins. **A)** Purified integrases (C3, C6, C7) were incubated with indicated plasmid DNA substrates and activity determined by real-time PCR using the primer 3 + 4 pair. Reactions comprised 17nM integrase and 1.3nM respective DNA substrate. n=2 ± SD. **B)** As in A, highlighting relative activities on *att*B x *att*BL substrate. **C)** PCR products from **A** resolved on agarose gel. Arrow indicates position of expected band indicating recombination. Repeat gel shown in Figure S3A.

**Figure 8.**
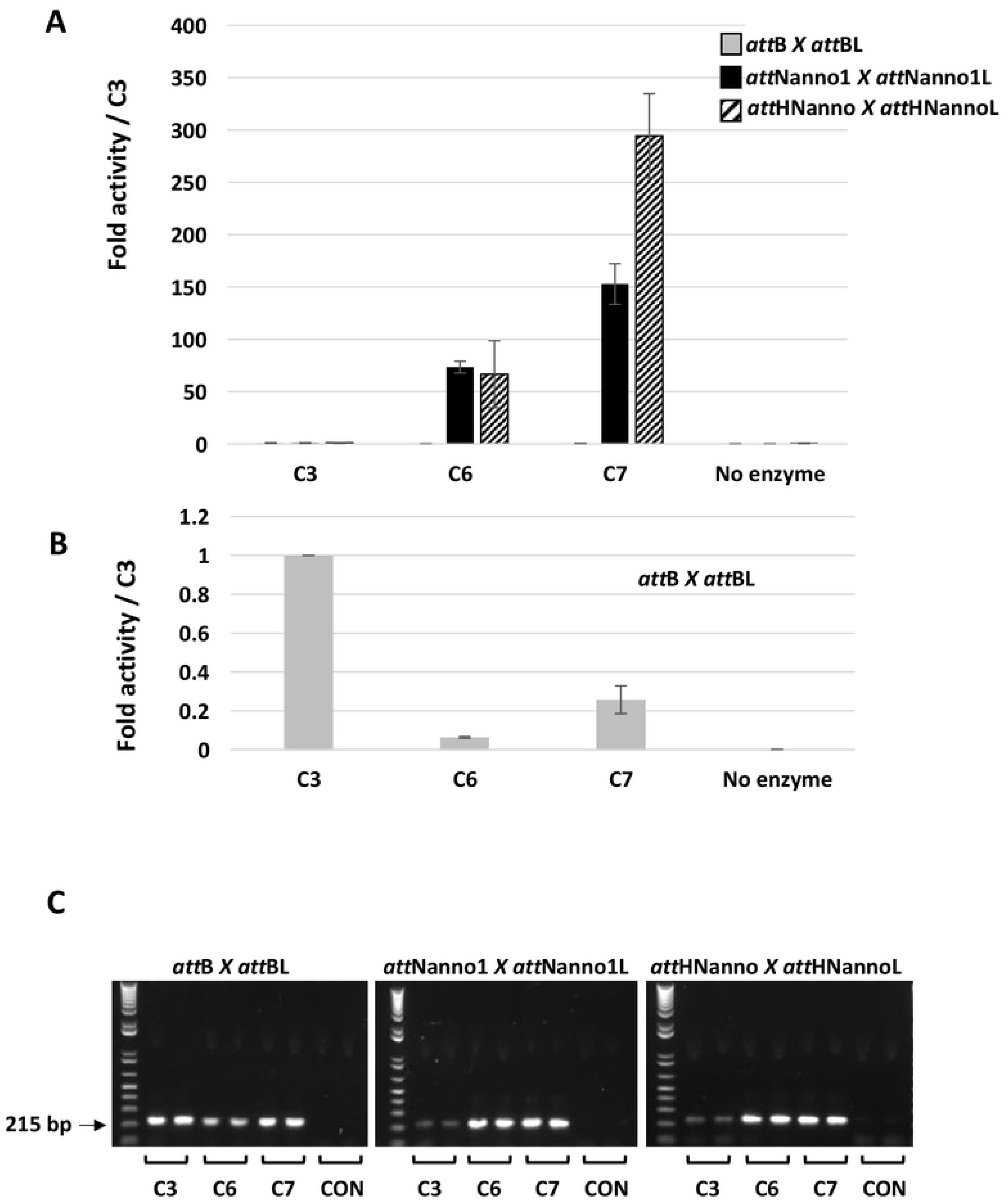
Recombination activity of selectants using purified proteins. **A)** Purified integrases (C3, C6, C7) were incubated with indicated plasmid DNA substrates and activity determined by real-time PCR using the primer 3 + 4 pair. Reactions comprised 1.7nM integrase and 0.13nM respective DNA substrate. n=2 ± SD. **B)** As in A, highlighting relative activities on *att*B x *att*BL substrate. **C)** PCR products from A resolved on agarose gel. Arrow indicates position of expected band indicating recombination. Repeat gel shown in Figure S3B.

### Molecular Dynamics Simulations

The roles of the selected N99D and H329R mutations were further investigated using molecular dynamics (MD) simulations. Structures of the wild type and mutant Int tetramer – DNA complexes remained stable throughout the simulations. For wild type Int bound to *att*B substrate DNA (*attB*-Int complex), the side chain of N99 interacts with the amide of guanine (DG8) for ∼95% of the simulation (Figure 9, S4). Note that DG8 base pairs to the *att*B cytosine boxed in Table 1. Replacement of N99 to D99 (*att*B-Int^C7^ and *att*B-Int^C3_N99D^ complexes), results in clear loss of the interaction with DG8, possibly explaining lack of activity of variants comprising N99D mutation on *att*B substrate (Figure 9, S4, S5A). The side chain of H329 was found to interact with D351 from the neighboring chain in the tetramer for ∼25% of the simulation time in the *att*B-Int complex. However, the H329R mutation resulted in an R329 – D351 salt bridge interaction that was stable for 60% of the simulation time for the *att*B-Int^C3_H329R^ complex (Figure S4, S5B). No difference was observed between the N99-DG8 interaction which remained stable as in the *att*B-Int complex. The combined H329R and N99D mutations present in C7 therefore result in loss of the N99-DG8 interaction but improved R329-D351 interaction (*att*B-Int^C7^ complex, Figure 9, S4, S5C). When DG8 in *attB* is replaced with cytosine (DC8) as present in *att*Nanno, no interaction is observed between this base and N99 of Int (*att*B*^DC8^*-Int complex, Figure 9, S5D). Note that DC8 base pairs with the guanine residue present in the variant *att* sequences (boxed in Table 1). Furthermore, the H329-D351 interaction was reduced from 25 % to only 10 % of simulation time in this complex (Figure S4). Similar interactions were observed for *att*B*^DC8^*-Int^C3^ complex simulations, again shedding light on the poor activity of Int/Int-C3 on the variant *att* sequences. Replacement of N99 with D99 results in formation of a stable network of interactions between the λ integrase with bound DNA in the *att*B*^DC8^*-Int^C3_N99D^ and *att*B*^DC8^*-Int^C7^ complexes (Figure 9, S4). This network engages D99 in interactions with both the amine of cytosine (DC8) and the K95 sidechain through formation of a salt bridge. Additional interactions are observed between the sidechain of K95 and the amine of adenine (DA21), the amide of guanine (DG22) and the nitrogen N3 of adenine (DA21). This large network of interactions was observed to be stable for > 75% of the simulation time and is likely a key determinant of the observed specificity switch. In addition to these interactions, the R329 – D351 interaction was observed for ∼ 60% of the simulation time in the *att*B*^DC8^*-Int^C7^ complex (Figure S4, S5E). No significant differences were observed for the R329 – D351 interaction in the *att*B*^DC8^*-Int^C3_H329R^ complex compared to *att*B-Int^C3_H329R^ (Figure S5F).

**Figure 9.**
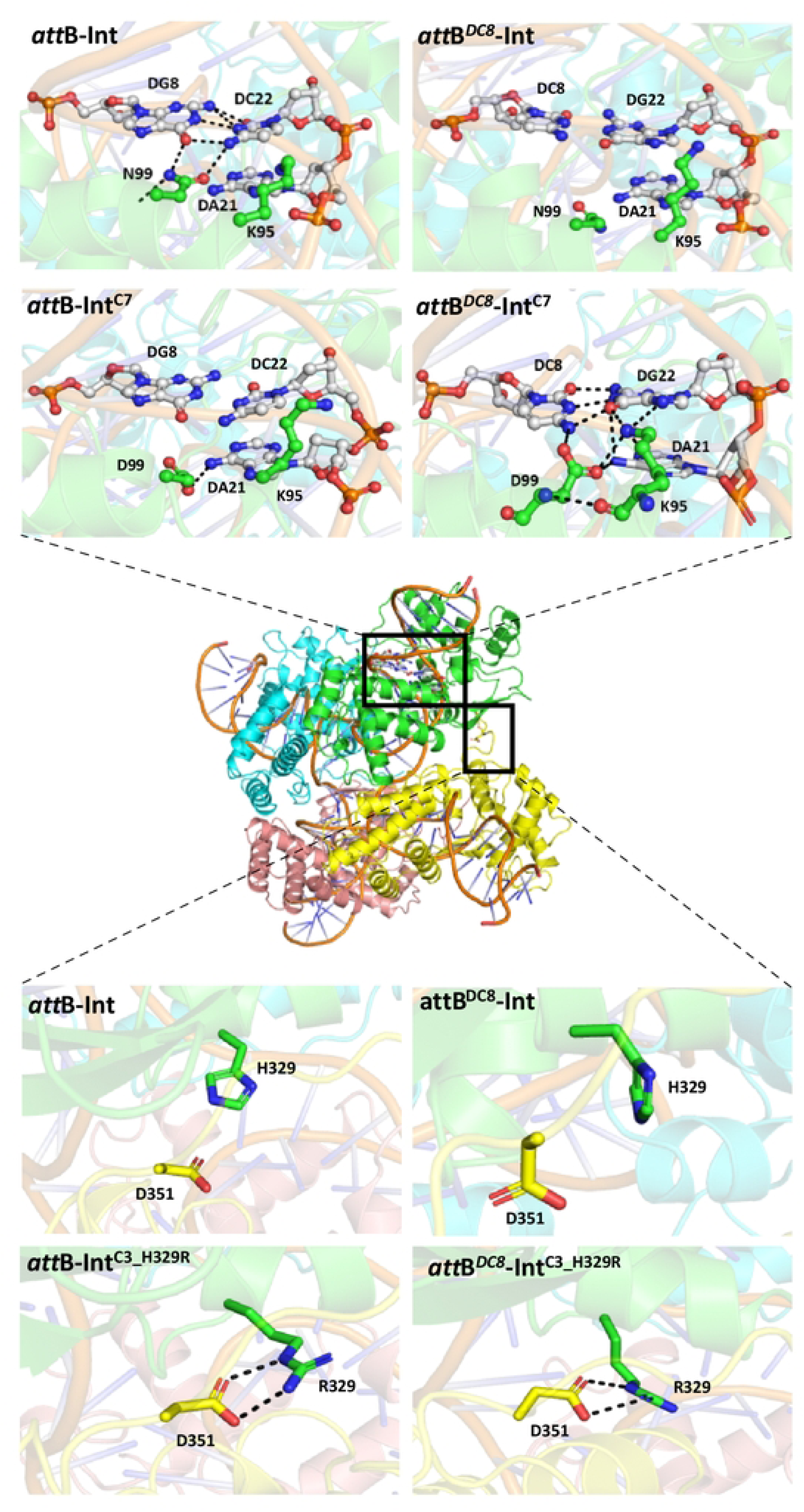
Molecular dynamics simulations of Int – DNA complexes. Tetrameric structure of Int shown in center with individual chains coloured differently. Boxed regions are expanded in the images above to show interactions of Int and Int C7 with *att*B or the variant *att*B*^D^*^C8^. Images below depict intra-chain interactions within Int and Int^C3_H329R^.

### In situ targeting of genomic att*H*Nanno by Int-C7 resulting in delivery of functional payloads

Int-C7 was next assayed for intermolecular recombination of a circular target vector carrying *att*HNannoL into the genomic 21bp *att*HNanno site on human chromosome 11. We generated a eukaryotic expression vector for Int-C7, termed pEF-ss-Int-C7-CNLS, coding for Int-C7 with a C-terminal nuclear localization signal (NLS) derived from the Simian Virus 40 (SV40) large T-antigen. Transfection into either HEK293ft or HT1080 cells following by Western blotting indicated stable Int C7 NLS expression at levels comparable to Int C3-NLS (Figure 10).

**Figure 10.**
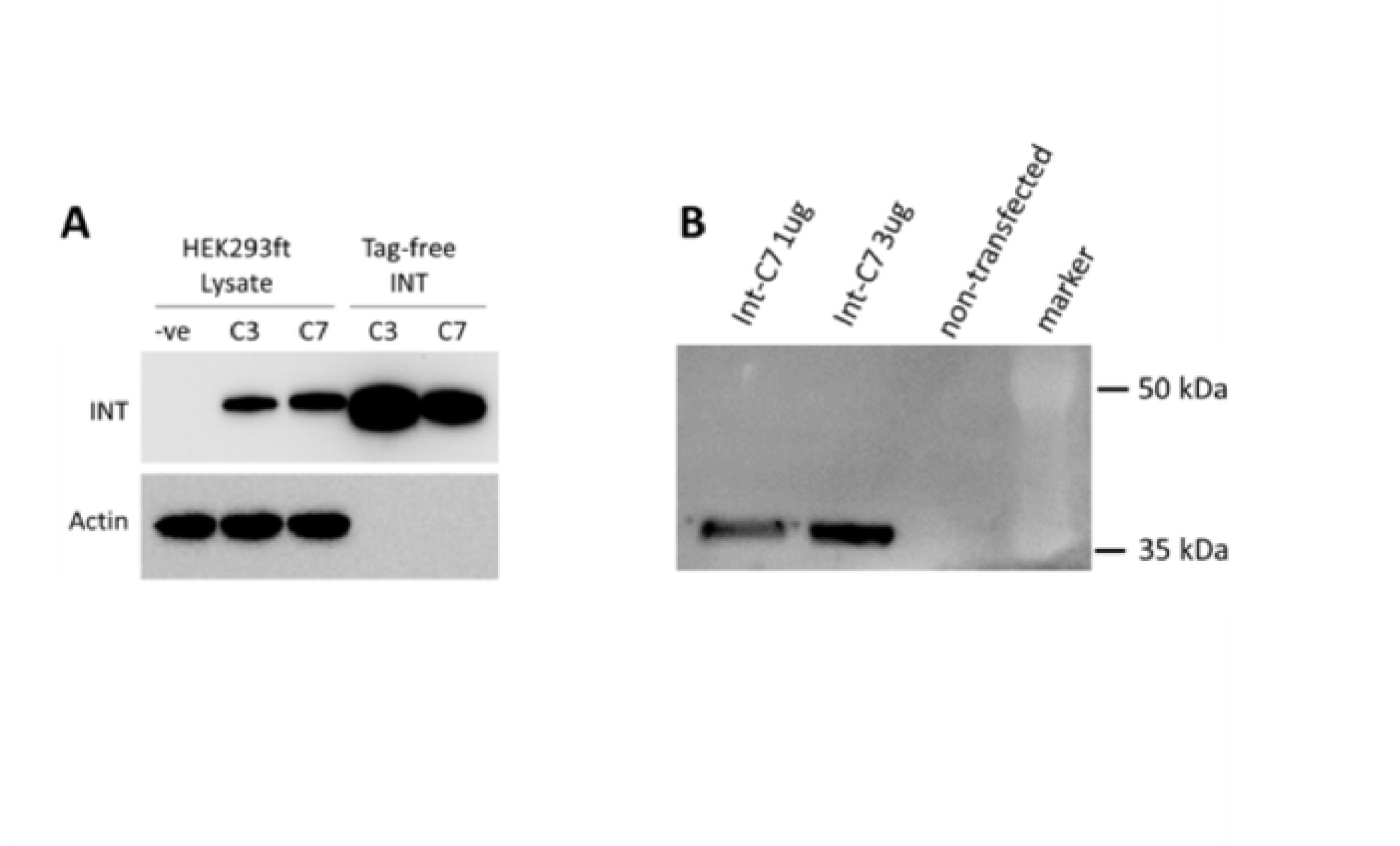
Western blot analysis of expressed integrase variants in human cells. **A)** Expression vectors for Int-C3 and Int-C7 were transfected into HEK293ft cells, and cell lysates were prepared by standard methods and analyzed by Western blotting using mouse anti Int-C3 antibodies. Both integrase variants are stably expressed, exhibiting slightly increased molecular weights due to the presence of a C-terminal nuclear localization signal (NLS) when compared with the corresponding purified proteins (from *E.coli*) loaded as controls, which do not have a tag. Endogenous actin protein was probed to control for loading of lysates. **B)** Two different amounts of expression vector for Int C7, as indicated, were transfected into human fibrosarcoma HT1080 cells and analyzed by WB as in (A), with lysate from non-transfected cells serving as negative control.

In order to target the genomic *att*HNanno sequence for integration *in situ,* we chose the following strategy (Figure 11): A 6.5 kb target vector carrying *att*HNannoL plus a selection marker and a GFP expression cassette as payload was co-transfected with expression vectors for Int-C7 and co-factor scIHF2 (40) into human HT1080 cells, followed by puromycin selection at 48 hrs. Post-selection, genomic DNA from bulk selectant cultures was isolated and PCR employed to identify the left and right recombination junctions (Figure 12). The results showed that with 750ng and 1000ng of transfected target vector, both junctions can be generated by nested PCR, which was verified by PCR product sequencing (Figure 12).

**Figure 11.**
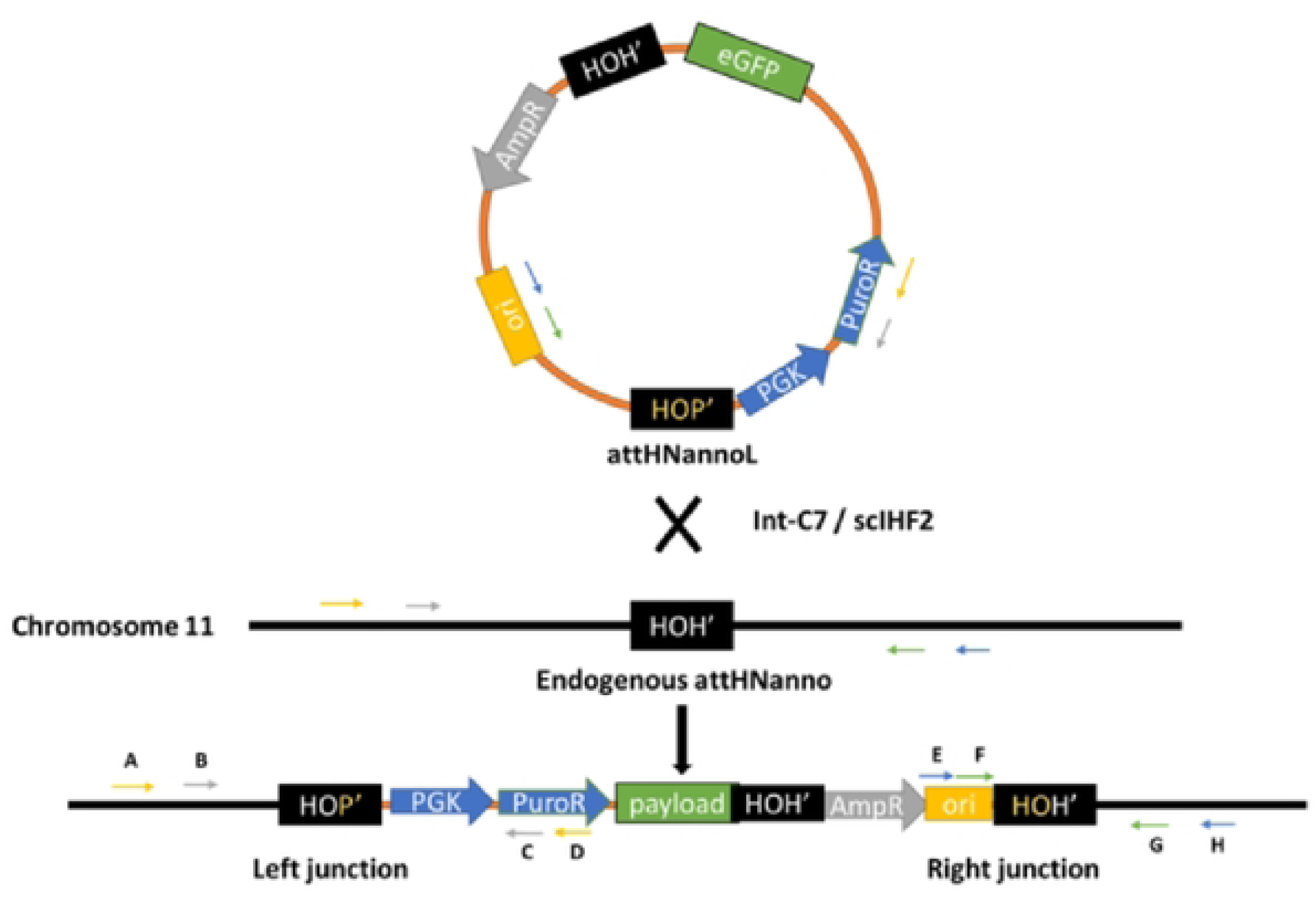
Targeting strategy for the genomic *att*HNanno sequence on chromosome 11. See text for details. The primers used to detect successful intermolecular recombination events at the junctions are indicated.

**Figure 12.**
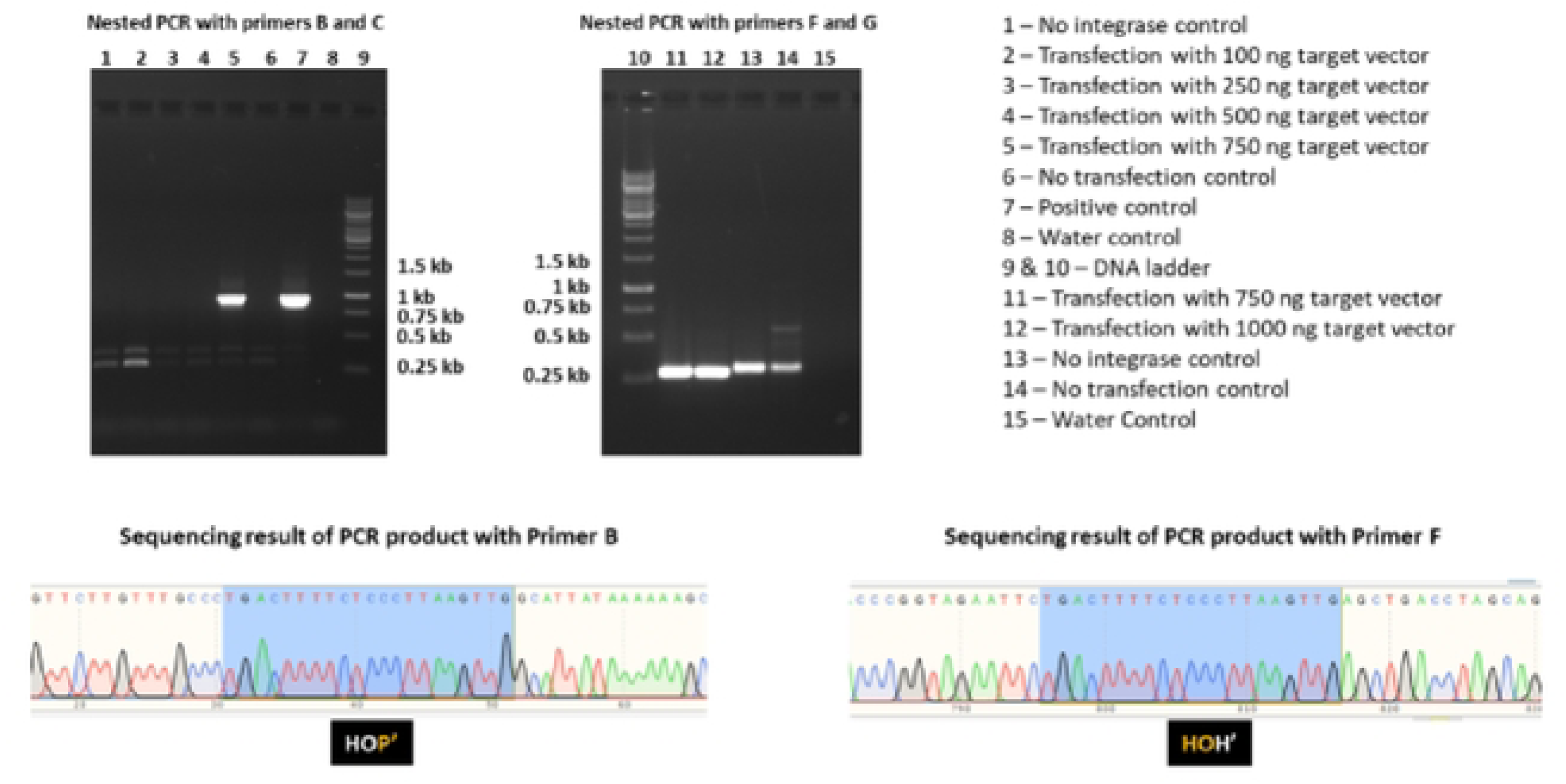
Detection of recombination events in bulk selectants using PCR. See Figure 11 for corresponding positions of PCR primers. The sequencing results of products for left and right junction are shown at the bottom.

In this experimental set-up, intramolecular recombination *in situ* between *att*HNanno x *att*HNannoL on the target vector (Figure 11) can first result in a seamless vector carrying only *att*HNannoL, which then recombines into the locus on chromosome 11. However, PCR/sequencing analysis of the junctions revealed that unrecombined plasmid DNA has been inserted by Int-C7 into the genomic locus. This indicates that contrary to expectations on thermodynamic grounds, Int-C7 surprisingly catalyzed intermolecular recombination more efficiently than intramolecular recombination, at least under the chosen experimental condition and with this substrate.

We screened individual cell clones from co-transfections and identified positive lines such as CL6, which showed the predicted junction product in the primary PCR as exemplified in Figure 12. Southern analysis indicated single copy payload integration into *att*HNanno in 3 out of 4 CL6 subclones (Figure 13A, B). The identity of the products was verified by sequencing. Furthermore, transgene expression (GFP) from this targeted human locus was homogeneous and sustained in the three CL6 subclones, which could be important for future gene/cell therapy approaches (Figure 13C). Using the same strategy, we could also demonstrate successful targeting of the endogenous *att*HNanno site in bulk selectants of HEK293ExPi cells (not shown).

**Figure 13.**
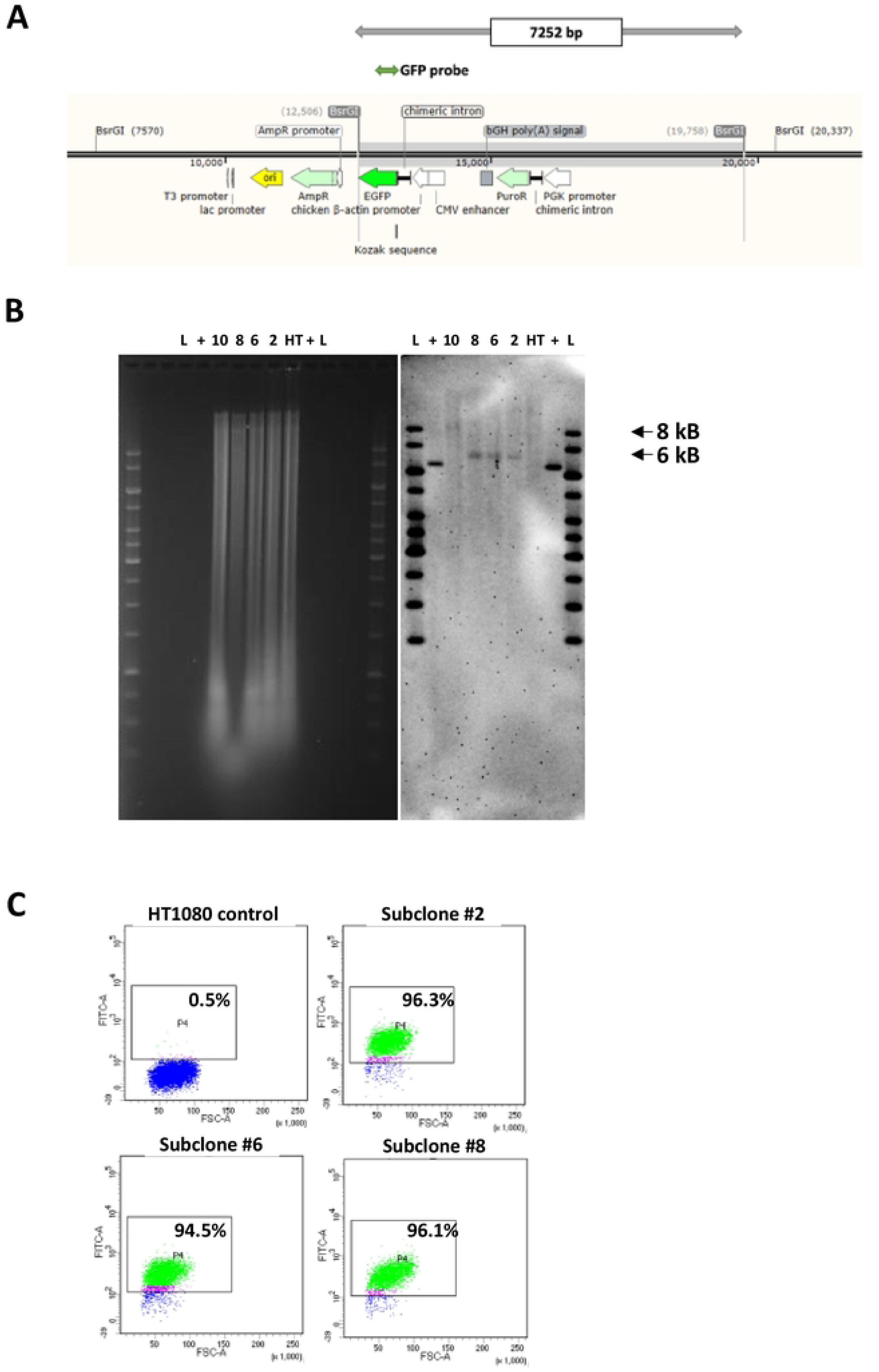
Single-copy targeted insertion of DNA payload into HT1080 *att*HNanno site. **A)** Feature map highlighting restriction sites and predicted orientation of GFP-expressing payload when integrated into *att*HNanno site. **B)** Southern blot indicating single-copy insertion of payload in subclones 2,6 and 8 determined by presence of expected 7.3 kB band (highlighted in **A**). ‘+’ corresponds to positive control DNA **C**. FACS analysis of HT1080 subclones 2, 6, and 8 showing increased GFP expression over control cells.

Bioinformatics analysis of the genomic *att*HNanno locus (chr11:110,828,911) shows the target sequence to be located in a gene-deserted region, with a single exon of pseudogene HNRPA1P60 positioned about 40 kb upstream (Figure 14A). No enhancer regulatory elements appear to be present in the vicinity of this locus. Furthermore, in the human pluripotent embryonic stem cell line H1, this locus belongs to a topologically associated domain (TAD) (Figure 14B), with an epigenetic H4K20me1 mark about 1.5 kb downstream (Figure 14C). Taken together, these findings indicated that the locus can be considered a safe harbor site favorable for transgene expression.

**Figure 14.**
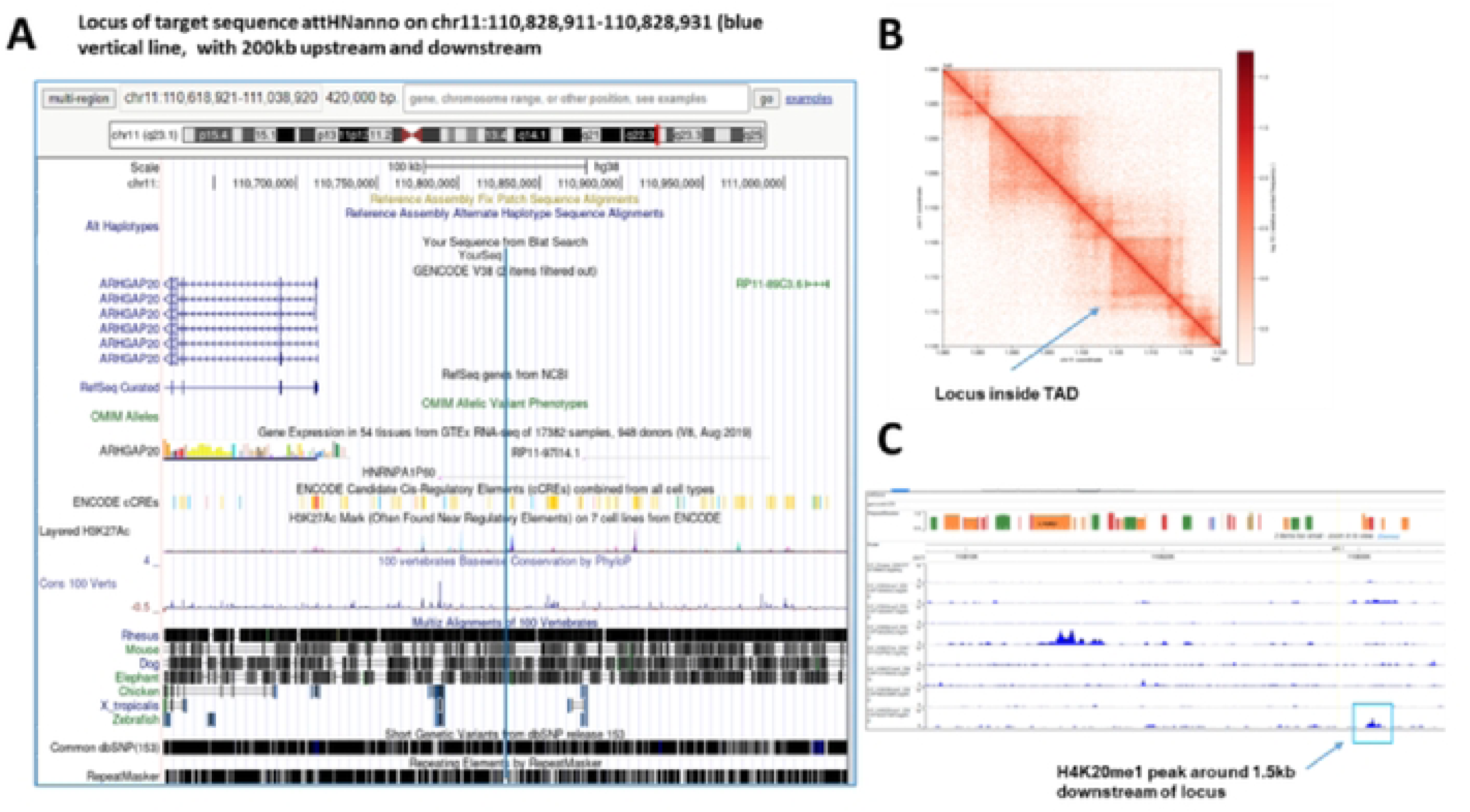
Bioinformatic interrogation of the *att*HNanno locus. **A)** Analysis of the genomic *att*HNanno locus (chr11:110,828,911) shows the target sequence to be located in a gene-deserted region. **B)** No enhancer regulatory elements appear to be present in the vicinity of this locus. **C)** An epigenetic H4K20me1 mark is located ∼ 1.5 kb downstream.

## Discussion

We have described selection of λ integrase variants displaying notable specificity switches towards pseudosites identified in microalgae and human genomes. Attempts at targeting the *att*Nanno1 and *att*Phae2 pseudosites *in vivo* in the host microalgae using the engineered Int variants have thus far been unsuccessful, most likely due to problems related to the efficiency of both foreign DNA uptake and transient expression of transgenes. The novel *att*HNanno site that we identified in mammalian cells was, however, successfully targeted. Bioinformatic analysis suggests *att*HNanno is a safe harbor that can now be targeted by the orthogonal Int C7 enzyme.

Residues N99 and H329 of λ integrase have previously been substituted for the corresponding amino acids (D and R respectively) in the highly related bacteriophage HK022 integrase, enabling a specificity switch towards the HK022 *att*B site(38, 39). Notably, the *attB* site targeted by HK022 shares the cytosine to guanine transversion present in the algal and human sequences targeted in this study (Table 1). Identical selection of these amino acids in this study both further confirms their role in target site discrimination and illustrates how readily directed evolution can recapitulate natural selection. MD simulations provide additional insight into the orthogonal phenotypes observed in this study. A favourable network of interactions promotes binding and activity of Int variants comprising N99D in the DNA core binding domain to substrates with the cytosine to guanine transversion. H329 in the Int catalytic domain is distal to bound DNA. Mutation to R329 potentially introduces up to 4 intra-subunit salt bridges that could stabilize tetramer binding to DNA and favour recombination of non-cognate sequences. Future structural studies will shed light on the roles of these amino acids in substrate discrimination.

Scarless gene curing can be achieved by sequential recombinase mediated cassette exchange (RMCE)(41, 42). Diseased genes flanked by pseudo *att* sites recognized by Int are excised in a first reaction and then replaced with the correct wild-type gene in a second reaction. RMCE is limited by the availability of pseudo sites and cognate enzymes targeting these. The highly active C7 Int variant along with its orthogonal *att* site potentially raises the number of genes that can be targeted. The odds of identifying appropriate pseudo-sites will also be improved by searching for flanking motifs orthogonal to C3 and C7 and subsequent combined use of these enzymes in a dual RMCE reaction.

In conclusion, we have evolved highly active and orthogonal Int variants with potential applications in microbial and human genome engineering.

## Acknowledgements

We thank Dr Oh Zhen Guo for blast sequence searches. This work was supported through a grant from the National Research Foundation-Competitive Research Programme, Singapore to PD (NRF-CRP21-2018-0002).

